# Task-uninformative visual stimuli improve auditory spatial discrimination: the ambiguous contribution of relative reliability

**DOI:** 10.1101/2022.08.24.505112

**Authors:** Madeline S. Cappelloni, Sabyasachi Shivkumar, Ralf M. Haefner, Ross K. Maddox

## Abstract

The brain combines information from multiple sensory modalities to interpret the environment. These processes, collectively known as multisensory integration, have been modeled as Bayesian causal inference, which proposes that perception involves the combination of information from different sensory modalities based on their reliability and their likelihood of stemming from the same causes in the outside world. Bayesian causal inference has explained a variety of multisensory effects in simple tasks but is largely untested in complex sensory scenes where multisensory integration can provide the most benefit. Recently, we presented data challenging the ideal Bayesian model from a new auditory spatial discrimination task in which spatially aligned visual stimuli improve performance despite providing no information about the correct response. Here, we tested the hypothesis that, despite deviating from the ideal observer, the influence of task-uninformative stimuli was still dependent on the reliabilities of auditory and visual cues. We reasoned shorter stimulus durations should lead to less reliable auditory spatial encoding, and hence stronger effects of more reliable visual cues, which are easily localized even at short durations. While our results replicated the effect from our previous study across a wide range of stimulus durations, we did not find a significant increase in effect size with shorter stimuli, leaving our principal question not fully answered.

## INTRODUCTION

When we observe our world, we must parse information that originates from many sources and is encoded through multiple sensory modalities. If we are to accurately perceive the world around us, especially when our environment is complex, we must decide how sensory cues are related, which ones are important, and how reliable they may be. The way our brain integrates these cues, especially across modalities, has a drastic effect on the resulting perception. For example, we can easily understand that seeing a violin and hearing a violin’s sound allows us to identify that a violin is playing. Less obviously, when listening to a concert with several musicians playing together, seeing the violin player’s bow movements may help you identify the notes of the melody. The rules governing the combination of sensory cues aren’t fully known and have typically been studied under a fairly limited range of circumstances that are closely related to the former scenario. The current work investigates situations akin to the latter scenario to explore how integration in complex environments can help us perceive and process our world.

Bayesian models like cue integration (Ernst & Banks, 2002) and more recently causal inference (Körding *et al.*, 2007) have formally described an optimal strategy for combining cues in complex scenes. In these models, each cue is treated as a measurement of the stimulus with a Gaussian likelihood of the stimulus based on that measurement. The multisensory measurement is then a combination of unisensory measurements weighted by the inverse of their relative variances, such that a narrower likelihood, indicating more reliable information, will have more influence on the combined percept. In the classic cue integration model *all* stimuli are combined in this way, but causal inference adds another inference layer to the model, in which the degree of cue integration depends on the probability that both measurements actually arose from the same event in the world (Körding *et al.*, 2007).

The causal inference model ultimately separates the issue of parsing complex information into two steps. The first step is the inference over causes and concerns potential relationships of stimulus cues in a scene. The second step is the combination of those cues resulting from the same cause by accounting for the reliability of the information they provide. We point out that for a given task there are two types of potentially helpful sensory information in a scene: 1) task-informative cues that provide information about the correct perceptual decision, 2) scene-relevant cues that may help the observer parse the scene, but do not give any information about the correct decision. For example, when listening to multiple people speak in a crowded room, knowing a person’s location offers no information about the content of their speech, and is thus task-uninformative, but helps a human observer to parse the overall scene (making it scene-relevant). While an ideal observer’s performance would not depend on such a cue, we have shown in prior work that human observers benefit from both task-informative and scene-relevant (but task-uninformative) cues (Cappelloni *et al.*, 2019).

Specifically, we engaged observers in a new task to test for the influence of task-uninformative cues on perception in humans and found a multisensory effect of task-uninformative visual stimuli on auditory spatial processing (Cappelloni *et al.*, 2019). We asked listeners to perform a concurrent auditory spatial discrimination task in which random visual stimuli were either spatially aligned with two symmetrically separated auditory stimuli or both located in the center of the screen, and we found a performance benefit when auditory and visual stimuli were spatially aligned. In both conditions, the visual stimuli did not provide any information about the correct decision in the task. The benefit provided by the spatially aligned visual stimuli is not explained by an ideal Bayesian observer nor can it be explained by endogenous/top-down attention (our very brief stimuli end before spatial attention could be redirected (Larson & Lee, 2013)). We showed analytically that the response of an ideal Bayesian observer did not depend on the task-uninformative visual stimulus in our experiments, suggesting that the improvement seen in real listeners may be the result of approximate rather than exact inference (Shivkumar *et al.*, 2022).

Unlike classical audiovisual experiments, our task provides an opportunity to both decouple task-informativeness from sensory reliability and to distinguish approximate from exact inference. Here, we engaged listeners in a variation of the auditory spatial discrimination task we used previously (Cappelloni *et al.*, 2019), this time modulating the reliability of auditory and visual stimuli by changing their duration. Because visual spatial information typically dominates audiovisual localization due to its better reliability (Witten & Knudsen, 2005) and a substantial amount of blur is required to noticeably decrease the relative reliability of visual stimuli (Alais & Burr, 2004), we hypothesized that shortening the duration of the stimuli would dramatically reduce auditory spatial reliability, but have minimal effect on visual spatial reliability. Furthermore, an effect of duration would be consistent with approximate inference, a topic that is explored in Shivkumar et al. (2022). We replicated the findings of our original experiment, which not only showed an effect of task-uninformative stimuli overall but also suggested that effect is larger for subjects with poorer auditory thresholds. Further, we showed a trend between auditory threshold and effect size across a population of subjects that is consistent with an effect of auditory reliability. However, the audiovisual effect did not differ significantly with duration.

## METHODS

**Figure 1.**
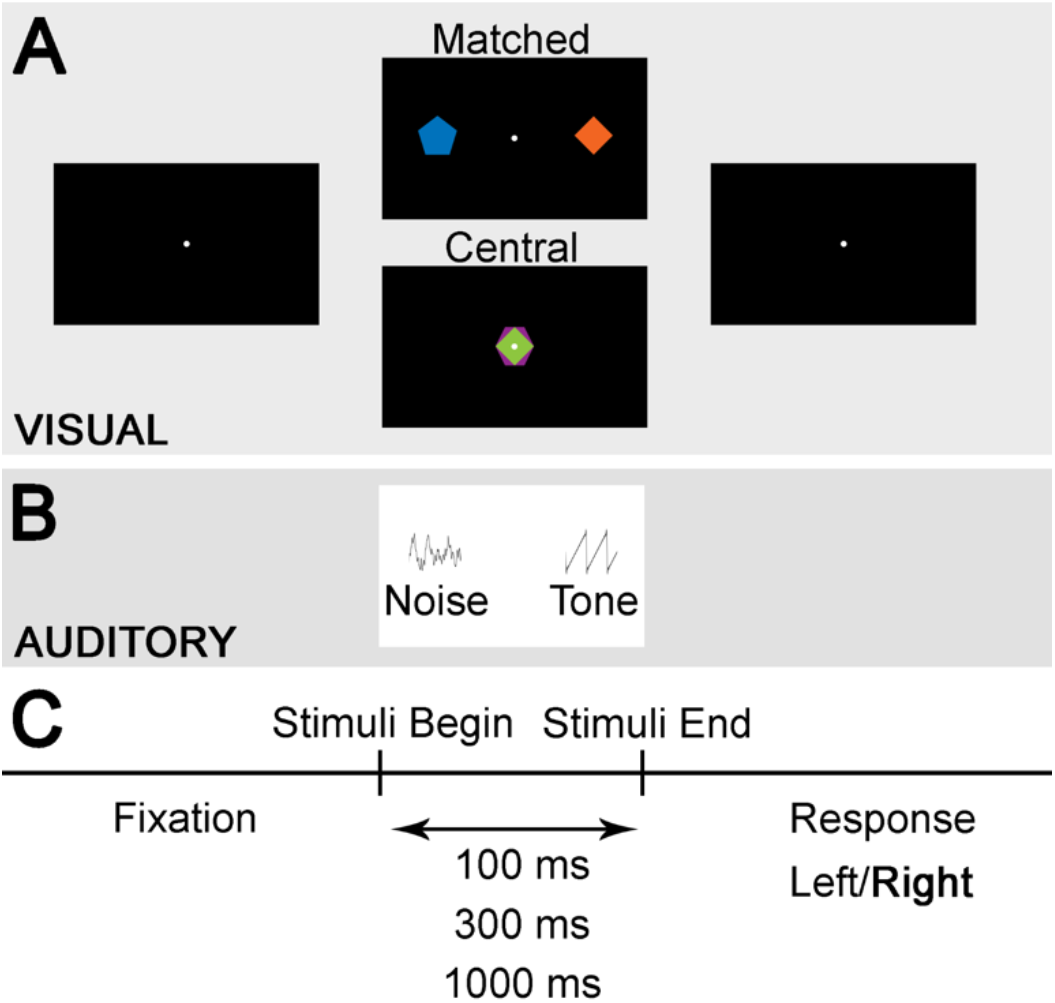
Demonstration of the task. A. Monitor views with frames representing each phase of the trial: during fixation, while the sounds are playing (depending on visual condition), and while the subject responds. B. Auditory stimuli begin after fixation and end before the response. C. Timeline of events from fixation through stimuli presentation to response.

### Participants

Participants (16 female, 4 male; ages ranging between 18 and 31, mean 21.5 +/- 3 years) with normal hearing (thresholds 20 dB HL or better at 500–8000 Hz) and normal vision (self-reported) gave written informed consent. They were compensated for the full duration of time spent in the lab. Research was performed in accordance with a protocol approved by the University of Rochester Research Subjects Review Board.

### Stimuli

Auditory stimuli were pink noise tokens and harmonic tone complexes with matching spectral envelopes, both bandlimited to 220–4000 Hz. Stimuli were generated and localized by HRTFs from the CIPIC library (Algazi *et al.*, 2001) using interpolation from python’s expyfun library as in Cappelloni et al. (2019), with the notable difference that here we generated the pink noise tokens and harmonic tone complexes to have three durations, 100 ms, 300 ms, and 1 s. Auditory stimuli were presented at a 24414 Hz sampling frequency and 65 dB SPL level from TDT hardware (Tucker Davis Technologies, Alachua, FL) over ER-2 insert earphones (Etymotic Research, Elk Grove Village, IL).

Visual stimuli were regular polygons of per-trial random color and number of sides. They were sized such that they could be inscribed within a 1.5° diameter circle. Colors were chosen to be clearly visible and have approximately uniform saturation and luminance, with the two stimuli in each trial having opposite hue as in (Cappelloni *et al.*, 2019). Any small discrepancies in perceptual luminance are randomly distributed across trials so as not to affect the results. Visual stimuli had the same onset and offset times as the auditory stimuli and thus matched their duration. To prevent overlap they were presented in alternating frames (Blaser *et al.*, 2000) on a monitor with a 144 Hz refresh rate.

### Task

Figure 1 shows an overview of the task. Each trial began when the subject fixated on a white dot in the center of the screen, confirmed with an eye tracking system (EyeLink 1000, SR Research). Then all four auditory and visual stimuli were presented concurrently for the duration of the trial (100 ms, 300 ms, or 1000 ms). After stimulus presentation, subjects were asked to respond with what side the tone was on by pressing a button. There were two visual conditions: one in which the visual stimuli were spatially aligned with the auditory stimuli and one in which the visual stimuli were collocated in the center of the screen.

We presented trials according to weighted one up one down adaptive tracks converging to 75% thresholds that adjusted the separation of the two sounds (Kaernbach, 1991). Separations were adjusted on a log scale such that separation increased by a factor of 2 when the participant responded incorrectly and decreased by a factor of 2^1/3^ when they responded correctly. By using log separation as the tracked variable, subjects are able to approach 0° separation with no danger of the sounds being actually collocated, and we follow the precedent of other adaptive tracking experiments regarding auditory space (Saberi, 1995). Each track had 130 trials and began at a starting separation of 10° azimuth. For each track, we randomized the number of trials with the tone on the left and right. There were six tracks, three durations by two visual conditions, that were randomly interleaved.

### Analysis

In order to obtain 75% thresholds we averaged the separation in log degrees at each reversal (skipping the first six reversals). We calculated threshold improvement as the difference between the separation thresholds of the two visual conditions (central – matched) in linear degrees. We also performed an ordinary least squares linear regression of the matched threshold to the central threshold data in log units and computed 95% confidence intervals of the regression coefficients to determine their significance.

Finally, we fit a generalized linear mixed effects model to the data. Using the glmer function from the lme4 package (Bates *et al.*, 2015) in RStudio (Version 1.4.11.06), we fit our model with a logit link function according to the following equation.

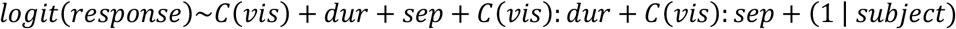

The model was fit to each trial, with the outcome measure being response (1 for correct, 0 for incorrect). Categorical visual condition (*C*(*vis*)), duration (*dur*), and separation (*sep*, in log units) were fixed effects as were the interactions of visual condition with duration and separation. Since the experiment was fairly simple, our only random effect was subject.

## RESULTS

**Figure 2.**
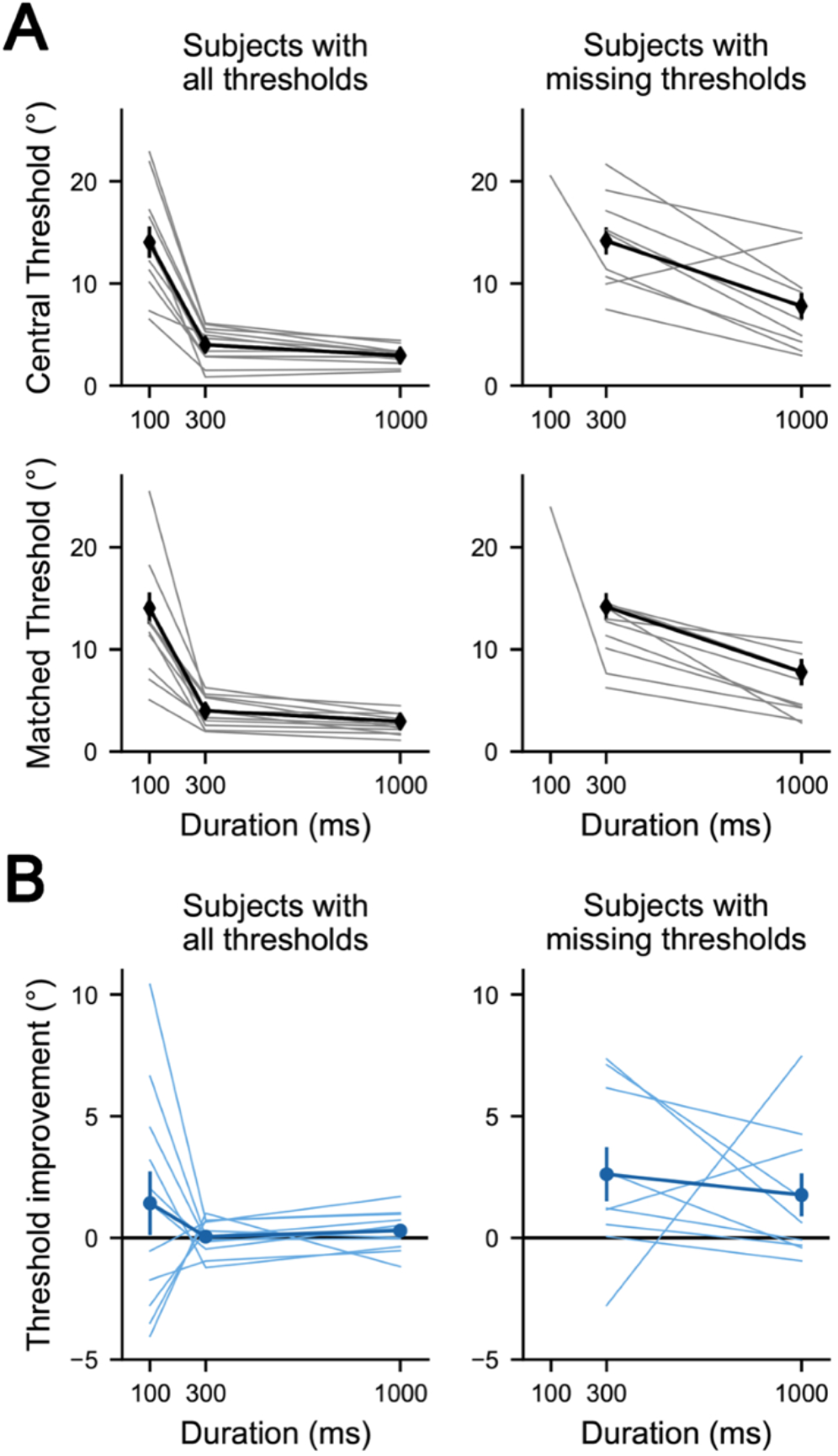
A. Thresholds for each duration in the central visual condition (top) and matched visual condition (bottom). Many subjects had missing thresholds (too large to measure accurately) in one or both visual conditions at 100 ms and are plotted in the right column (n=9) while the remainder are plotted on the left (n=11). B. Improvements in threshold for the two groups of subjects: those who could perform the task at all durations (left), those had one or both thresholds missing at 100 ms (right).

Subjects improved their task performance, indicated by a decrease in threshold, asymptotically in both visual conditions as the duration of the auditory stimuli increased; however, there was considerable variation in subject performance. Only 11 of 20 subjects were able to perform the auditory discrimination task at the shortest duration such that we could calculate a separation threshold (Figure 2A). Subjects in this group had a large decrease in threshold between 100 ms and 300 ms, but did not improve further for 1000 ms stimuli. For the remainder of the subjects who had thresholds too big to calculate in either or both 100 ms conditions, they improved their threshold between 300 ms and 1000 ms. In a linear mixed effect model of all subjects combined (Supplemental Table 1), only duration and separation (both p<2×10^-16^) had significant effects on performance. Neither visual condition nor the interactions of visual condition with separation and duration had significant effects on threshold (p=0.64, p=0.29, and p=0.42 respectively).

We defined “threshold improvement” as the linear difference between the central and matched visual conditions and used it to measure the size of the visual benefit (Figure 2B). Differences in individual auditory spatial processing ability indicate that auditory reliability was not uniform within a given duration condition.

**Figure 3.**
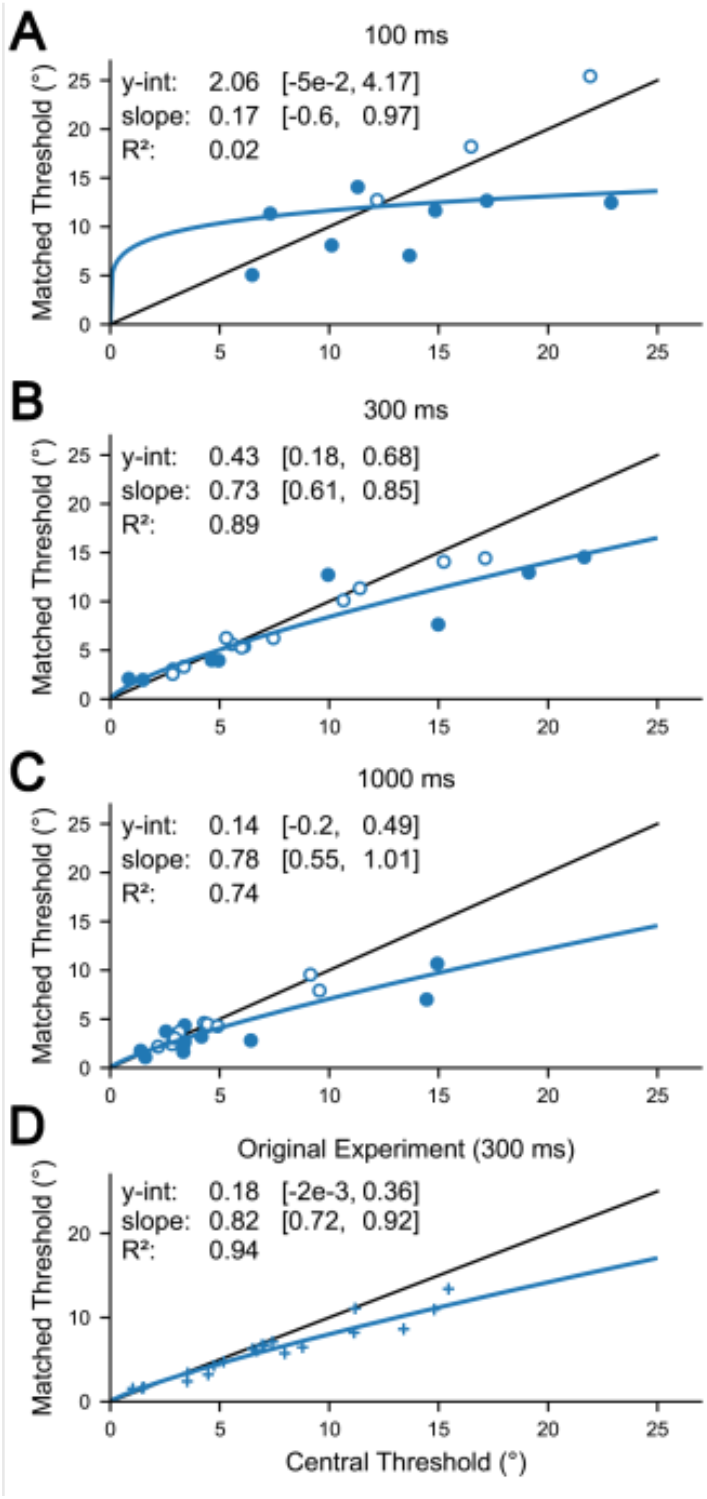
Linear regression of matched threshold against central threshold computed in log units and plotted in linear units. Parameters and their confidence intervals are expressed in log units. Solid markers indicated significant differences between the two visual conditions based on within subject variation (confidence interval does not include identity line with **α**=0.05 uncorrected). Open markers indicate no significant effect of visual stimuli. A–C. Separate regressions for each duration. D. Regression of data from our previous study (Cappelloni et al., 2019).

We compared the thresholds measured in the central condition to those in the matched conditions by fitting a linear model in the space of log thresholds (shown in Fig. 3 after mapping back to linear thresholds for interpretability). If the visual cues have no effect on the perceptual decisions, we expect the regression to predict a line with zero intercept and slope of 1 in log thresholds (and thus also linear thresholds). Any deviations from this hypothesis would manifest as a non-zero intercept or a non-unity slope in log thresholds, leading to a curve in linear units (a linear dependency in log units becomes a power-law in linear units with the slope in log thresholds determining the exponent in linear units and the intercept in log units corresponding to the logarithm of the scale in linear units, see Analysis Methods). In the 100 ms condition data were highly variable and the slope and intercept of the regression model were poorly constrained (R2 = 0.02). However, the 300 and 1000 ms conditions were well fit with R2 = 0.89, 0.74 respectively. For 300 ms, the slope of the regression line was significantly less than one (p<0.05), indicating that improvement increased as central thresholds worsened. In this condition there was also an intercept significantly above zero (95% confidence interval does not include zero). In the 1000 ms condition, data trended towards the same pattern, but the 95% confidence interval of the slope includes one and the 95% confidence interval of the intercept includes zero. Reanalyzing the data from the original experiment (Cappelloni *et al.*, 2019), we found the linear model in log units to also be a good fit with R2=0.94 and a slope that was significantly less than (p<0.05) and positive intercept that was not significantly different from zero (0.18, p> 0.05).

## DISCUSSION

Our data are consistent with, but do not provide evidence to support our hypothesis that relative reliability modulates the multisensory effect of task-uninformative but scene-relevant visual stimuli. Providing weak support for a role of reliability in this task, trends across subjects but within duration conditions indicate a stronger effect for subjects with worse auditory thresholds. Contrary to our starting hypothesis, we found no significant difference between the duration conditions.

Our work replicates our previous finding that task-uninformative but spatially aligned stimuli benefit auditory spatial discrimination, with the new insight that this effect is strongest where auditory thresholds are large. For subjects who had central thresholds above approximately 5° separation in azimuth, we observed visual benefits of similar size and pattern in this and our original experiment (Cappelloni *et al.*, 2019). We did not, however, observe an improvement across subjects in the 300 ms condition as we had in the original experiment. There are a few differences between the paradigms that may have contributed to this. In our previous study, the size of the visual effect is clearest when looking at the percent improvement at the central threshold (the performance gain at a set separation), which we could not measure here due to our use of adaptive tracks to estimate threshold. Additionally, the visual stimuli preceded auditory stimuli by 100 ms in the previous study, whereas in this study, their onsets were all concurrent. That the data in the 300 ms condition and our original experiment show very similar trends nonetheless suggests that the visual benefit is relatively robust to small audiovisual asynchronies and changes in the distribution of stimuli across space due to the adaptive tracks.

Although the experiment was designed with the goal of comparing different duration conditions, we found that the 300 ms condition was most informative. At 300 ms the task is neither too hard nor too easy for the population of subjects and we were able to fit a model that not only describes the data well but also has well constrained parameters. In contrast, thresholds in the 100 ms conditions were highly uncertain. The most obvious explanation for the variability is that the task was extremely difficult and we were able to collect less data than for other conditions. Another source of uncertainty could be that 100 ms approaches the temporal limits of the underlying neural processing, interfering with the multisensory effect we observe at 300 ms. In contrast, when the stimuli are 1000 ms in duration, the task becomes very easy, and most people have separation thresholds better than 5°. Thresholds at a 1000 ms duration cover a limited range and the model parameters were poorly constrained, with wide confidence intervals. If we had been able to sample more subjects with poor thresholds in this condition, we speculate that the slope would be significantly less than 1 and intercept significantly greater than 0, as they were for 300 ms.

Further obscuring the effect of stimulus duration, subjects could be divided roughly into two groups with two different patterns of thresholds. Subjects who could reliably perform the task at 100 ms improved their performance when the duration was extended to 300 ms. However, these subjects did not further improve when the stimuli duration was 1000 ms, suggesting that they reached ceiling performance between 300 ms and 1000 ms. In contrast, the remaining subjects improved when the stimulus duration extended from 300 ms to 1000 ms. Without considering effects in individual subjects rather than the population, for which we did not have enough data, we could not establish a robust relationship between changing stimulus duration and the visual benefit even though weak evidence suggested one may exist.

### Follow-up Experiment

Lack of support for our hypothesis motivated us to design an experiment that addressed several issues from the main experiment in hopes of showing a clear effect of reliability on the integration of task-uninformative stimuli. We moved to a small-n design with stimuli catered to each subject’s ability to address the two factors that made it difficult to compare duration conditions: 1) subjects’ auditory thresholds varied widely, making across subject comparisons difficult and 2) many trials were wasted because they were either too easy or too hard. We also added an auditory-only control condition to address the question of whether the multisensory effect of task-uninformative stimuli is truly driven by a benefit of spatially aligned stimuli or is due to a decrement when the stimuli are centrally located. Finally, hoping to maximize our chances of measuring a significant effect by making the stimuli more natural, we presented stimuli from loudspeakers at multiple locations (rather than using HRTFs) and a virtual reality headset, which placed the visual stimuli in a 3D environment rather than a flat computer monitor. Full Methods and Results for this follow-up experiment are presented in Supplemental Materials.

Despite targeting our follow-up experiment to remedy as many factors as possible that could have contributed to ambiguity in our main experiment, our data do not show an effect of duration on effect size. Worse, in this new experimental setup we failed to replicate the original effect of the task-uninformative visual stimuli on performance.

Collecting several sessions of data from each observer did not produce the reduction in variability that we had hoped for. Here the variability across subjects was replaced with large variability from session to session. Compounding this issue, duration thresholds were measured at the beginning of the first session and stimulus strengths were based on them for all remaining sessions (at least one subject showed clear effects of learning after the initial session). In hindsight, the relevance of these initial thresholds to later sessions is dubious, particularly as they seem to be overestimated in one subject (who showed clear effects of learning after the initial session) or so brief as to be as uninformative as the 100 ms condition in our main experiment (two subjects’ thresholds were even shorter than 100 ms).

Adding an auditory-only control condition did not clarify the nature of the original multisensory effect. Results were too variable to allow for a meaningful comparison between the control and the two visual conditions since this modification reduced the number of trials in each condition, and variability in our more natural experimental setup was substantially larger than in our original one.

Presenting stimuli with loudspeakers and VR substantially improved the realism of those stimuli and anecdotally reduced subject fatigue. Since we did not replicate the original multisensory effect, it is unclear whether this realism has any beneficial or detrimental influence on multisensory processing.

## CONCLUSION

Here we aimed to expand our understanding of the multisensory effect we measured in Cappelloni et al. (2019), having proved that it could not be explained by exact Bayesian causal inference. Specifically, we hoped to investigate whether this deviation was modulated by stimulus. While our results replicate our original findings in a highly controlled experimental context, we did not find a statistically significant effect of stimulus duration. Furthermore, when we changed the experimental paradigm to make it more naturalistic, at the expense of being less controlled, the variability of our data increased to an extent that made it impossible to measure the original effect.

## SUPPLEMENTAL MATERIALS

**Supplemental Table 1:**
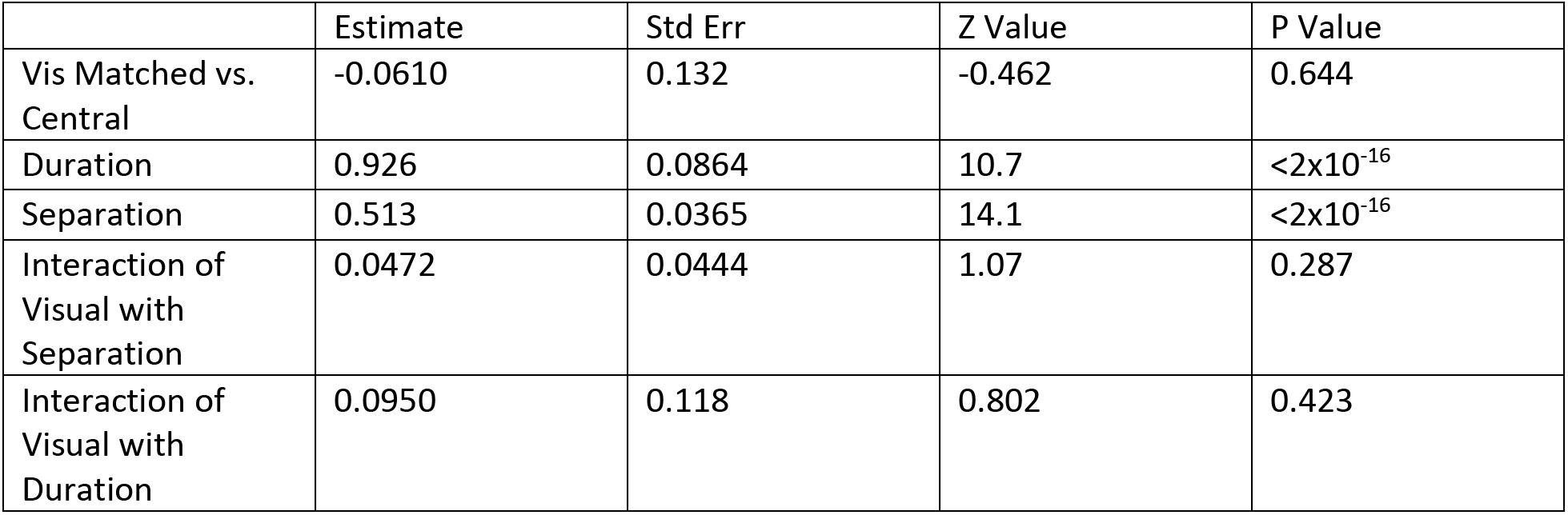
GLM for Experiment 1.

### Add-on Experiment Methods

#### Participants

Three participants (1 female, 2 male; of age 21–22) satisfied the same inclusion criteria from the main experiment. We originally planned a run of six subjects but stopped data collection when it became clear that even collecting more data would not clarify our results.

#### Equipment

All data collection took place in a large 6 m x 3 m (10 ft x 20 ft) soundproof booth. Visual stimuli were presented using an HTC Vive Proeye VR headset with a refresh rate of 90 fps. The virtual scene was a copy of the empty sound booth. We precisely calibrated the location of walls, light fixtures, and floor by capturing the location of three Vive 2.0 Trackers, averaging their x, y, and z coordinates to compare with a reference point, and applying a linear offset to all elements in the virtual scene. The Vive 2.0 Trackers are very accurate, with position errors on the order of only a few centimeters, and by using the average of three trackers we further improve our tracking accuracy. Auditory stimuli were presented with a loudspeaker array. The loudspeaker array contained of 53 KEF E301 speakers with 4° spacing (subtending a −104° to 104° range) in an arc with a radius of 2 m. The subject was seated in the center of the arc in an adjustable height chair such that their ears were roughly level with the center of the loudspeakers. Subjects held two HTC Vive Controllers, one in each hand, to report their responses.

**Figure 4.**
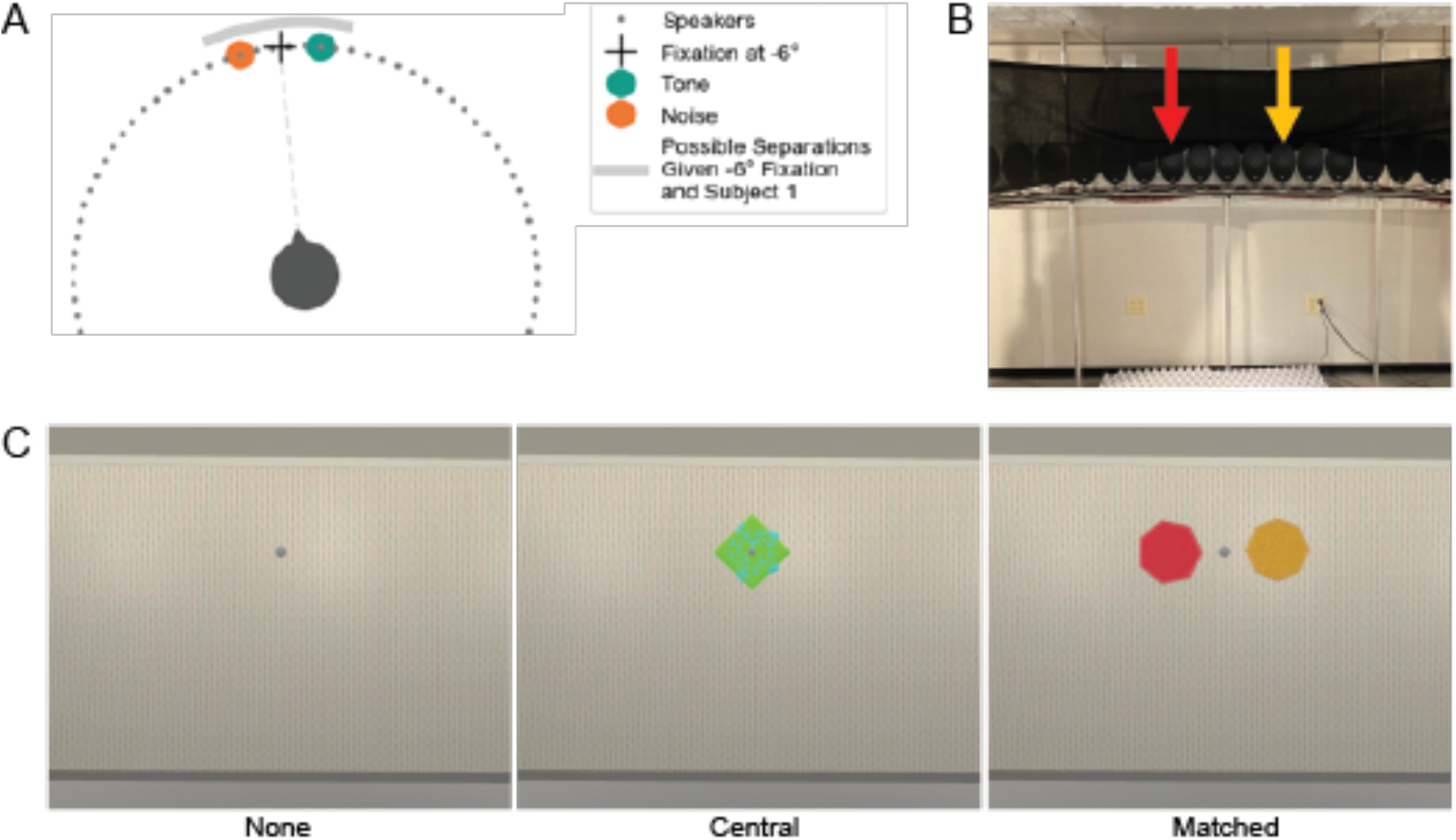
Setup for the booth. A. General layout. The subject sits in the center of the speaker array. An example trial is included with fixation at –6° and a 20° separation. Dashed line showed alignment of subject with the fixation that begins the trial. B. Speaker positions in the real booth. Colored arrows show a possible location of the two sounds with a central fixation and 16° separation. C. The three possible visual conditions given the configuration of sounds in B. The color alignment in the third panel is to emphasize the spatial connection between the real speakers and visual shapes, but the sounds are not associated with any color.

#### Stimuli

The same tone and pink noise stimuli were used from the main experiment. However, instead of localizing each sound by HRTF, the sounds were presented from a speaker at their true locations. Auditory stimuli were presented at a 48000 Hz sampling frequency and at 65 dB SPL.

Visual stimuli were gengons (regular polygons with extruded depth) of per-trial random number of sides (polygon face having between four and eight sides) and color. We slightly beveled the edges of the gengons so that their dimensionality was evident. Colors were chosen to have uniform saturation and luminance, with the two stimuli in each trial having their hue separated by 0.1 (instead of 0.5 as in the previous experiment) on a scale of 0 to 1. This deviation from the original experiment in color choice was due to the stimuli being more clearly distinguishable by their three-dimensional shape without an extreme color difference. Visual stimuli had the same onset and offset times as the auditory stimuli and thus matched their duration. To prevent complete occlusion of one stimulus by the other, both were given a depth texture such that an equal amount of each shape would occlude the other in a random pattern.

#### Task

Each trial began when the subject fixated on a grey sphere located at the center of each trial (randomly selected between −6° and 6° azimuth with 2° intervals). We verified fixation by determining that the normal forward vector of the VR headset was within a 2° tolerance of the fixation azimuth. Then, all stimuli were presented concurrently for the duration of the trial (customized per subject). After stimulus presentation, subjects were asked to respond with what side the tone was on by pressing a button on the VR controller in their corresponding hand. There were three visual conditions: one in which the visual stimuli were spatially aligned with the auditory stimuli, one in which the visual stimuli were collocated in the center of the screen, and one (not included in previous experiments) in which no visual stimuli other than the fixation sphere were present.

Subjects participated in four or five sessions of the experiment. In the first session, we determined the trial duration using weighted 1-up 1-down adaptive tracks converging to 70% thresholds (Kaernbach, 1991). Auditory stimuli were presented with a 20° separation in all thresholding trials. We randomly interleaved three tracks, one for each of the three visual conditions, and repeated each track three times for a total of nine threshold measurements. Trial duration increased after an incorrect response initially by 150 ms and then 60 ms after the first two reversals and decreased after a correct response initially by 50 ms and by 20 ms after the first two reversals. Each threshold was calculated as the average of the duration at reversals, skipping the first two reversals. The initial stimulus duration was 600 ms in the first set of tracks. In the second and third repetitions, we took the averaged threshold across the previous set of three tracks and added 150 ms. For the remainder of the experiment, we took the average of all nine tracks and used it as the shorter duration for that subject. The longer duration for each subject was twice that of the shorter duration. In the second half of the initial session, subjects completed half of a data collection session (for subject 1: 384 trials, for subjects 2 and 3: 396 trials).

For the remaining sessions, subjects completed approximately 800 trials. For subject 1, we tested a broader range of separations: 4° to 32° in 4° intervals. For subjects 2 and 3, we did not include the two largest separations so that we could increase the number of trials in each of the remaining conditions. Subject 1 completed 768 trials per session (3 visual conditions x 2 durations x 8 separations x 16 repetitions). Subjects 2 and 3 completed 792 trials per session (3 visual conditions x 2 durations x 6 separations x 22 repetitions). Trials from all conditions were randomly interleaved with right and left trials and fixation location counterbalanced. The first 10 trials of each data collection session (including the second half of the Initial session and all remaining sessions) were discarded as “warmup trials” and not included in any analyses.

#### Analysis

We used a two-step sigmoid fitting process to reduce the number of parameters in our threshold calculation, similarly to our methods from Cappelloni et al 2019. This involved an initial fit to establish slope and lapse rate across visual and duration conditions followed by a fit of midpoint (the separation where performance is the midpoint of chance and lapse rate) for each condition. Specifically, for each day we calculated the average percent correct for each separation (averaging across visual conditions, durations, and trials). The first fitting step was to fit a sigmoid to the average percent correct (across conditions) by separation (in log units, as in the main experiment) data. In the second fitting step, we fit individual sigmoids to data from each of six conditions (three visual x two duration) only allowing the midpoint parameter to change. We then calculated the 75% threshold (the separation at which the fit crossed 75% performance), the parameter of interest. Although we assume that lapse and slope parameters are consistent across visual and duration conditions, we do allow them to differ between sessions by repeating the entire fitting process for each day.

We fit a generalized linear mixed effects model with a logit link function using the glmer function from the lme4 package (Bates *et al.*, 2015). A separate model was fit for each subject, with every trial considered a separate data point. We modeled the following equation

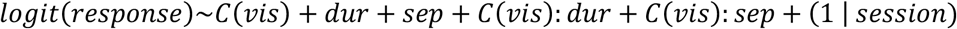

Where the outcome is whether the response was correct, and we consider fixed effects of categorical visual condition *(C(vis)),* duration *(dur),* separation *(sep,* in log units), interaction of visual condition with separation, and interaction of visual condition with duration. We also included a random effect of session (1 | *session).*

### Add on Experiment Results

**Supplemental Figure 1:**
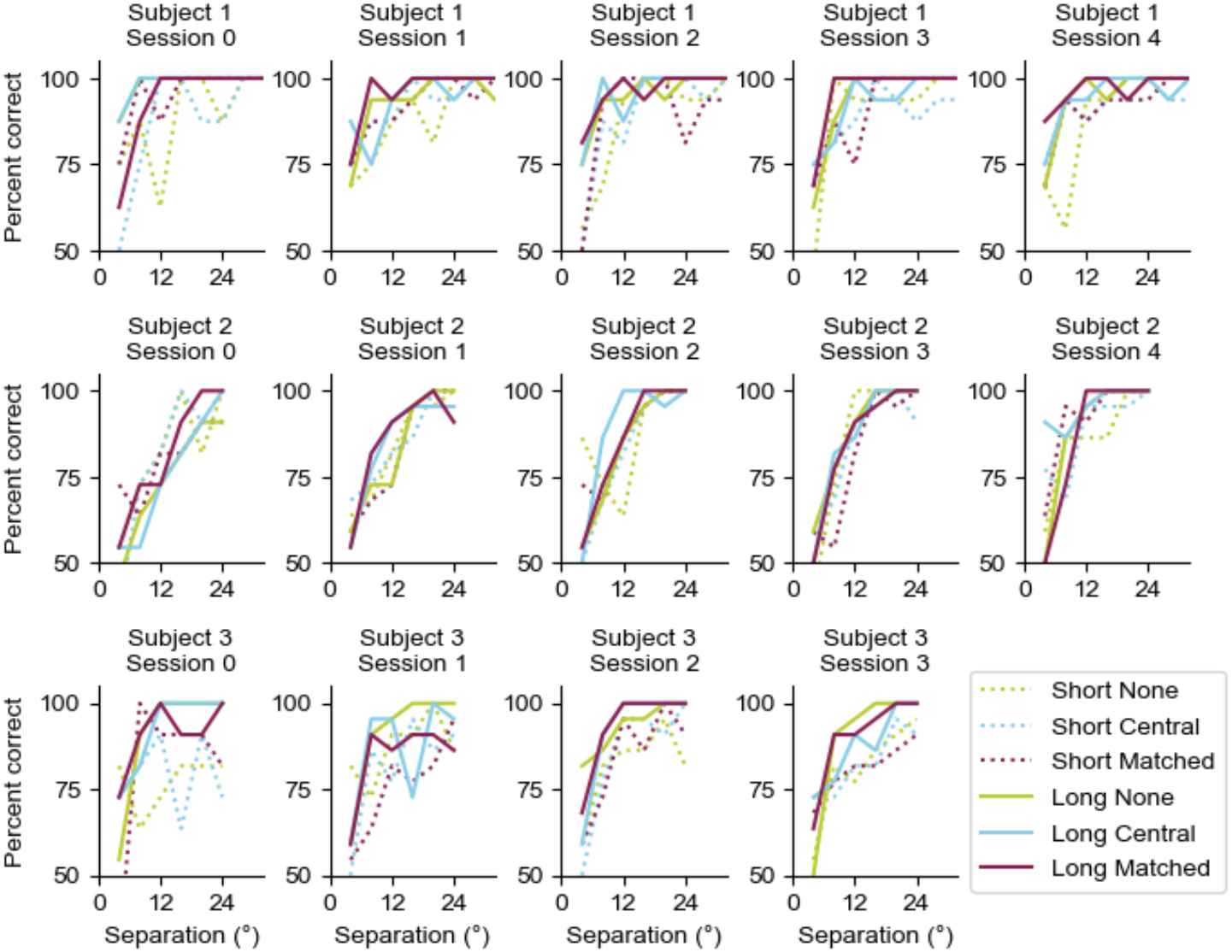
Experiment 2 data for each subject and session.

**Supplemental Figure 2:**
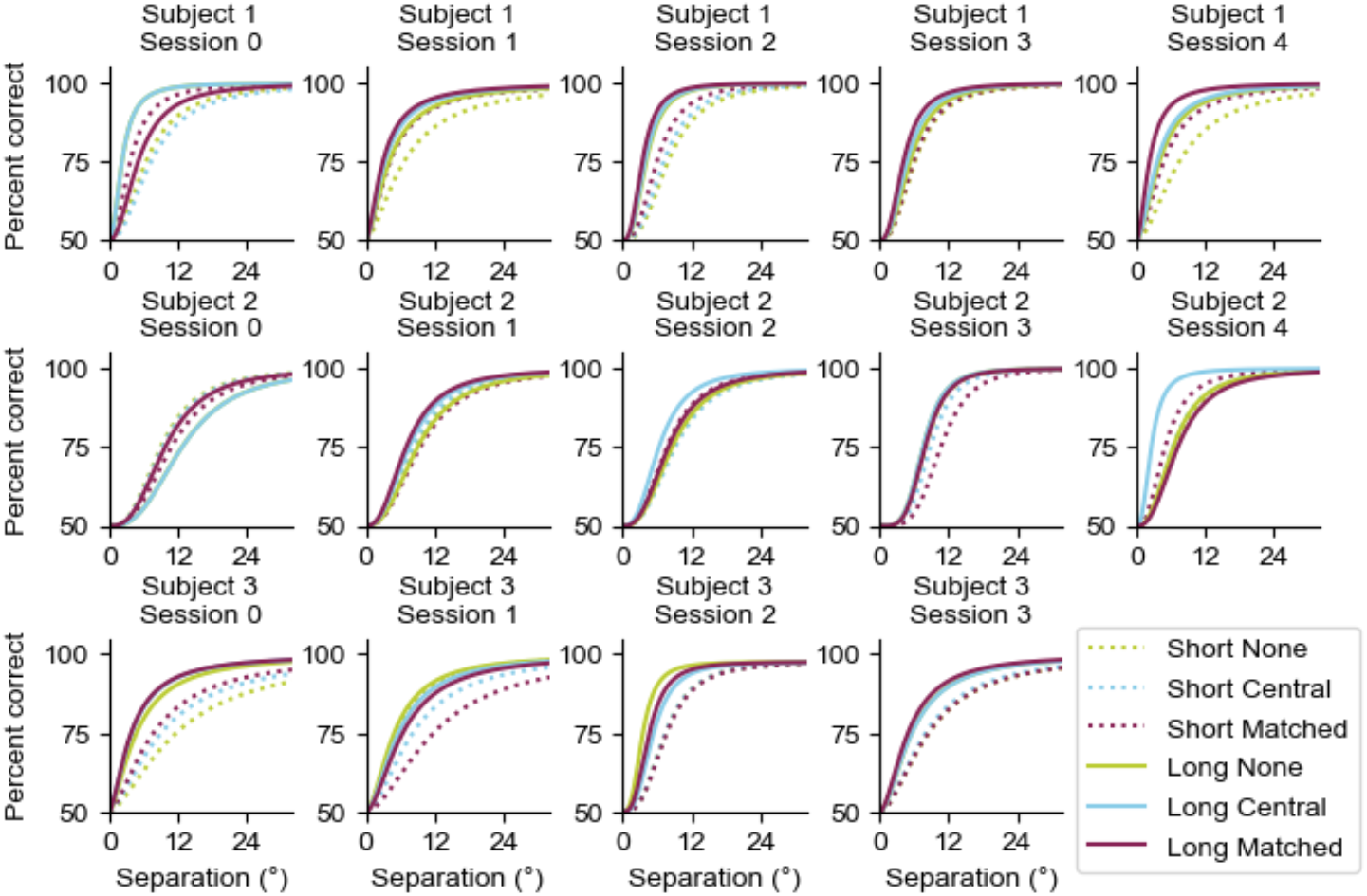
Experiment 2 fits for each subject and session. Error not shown due to its large magnitude obscuring the lines.

**Supplemental Figure 3.**
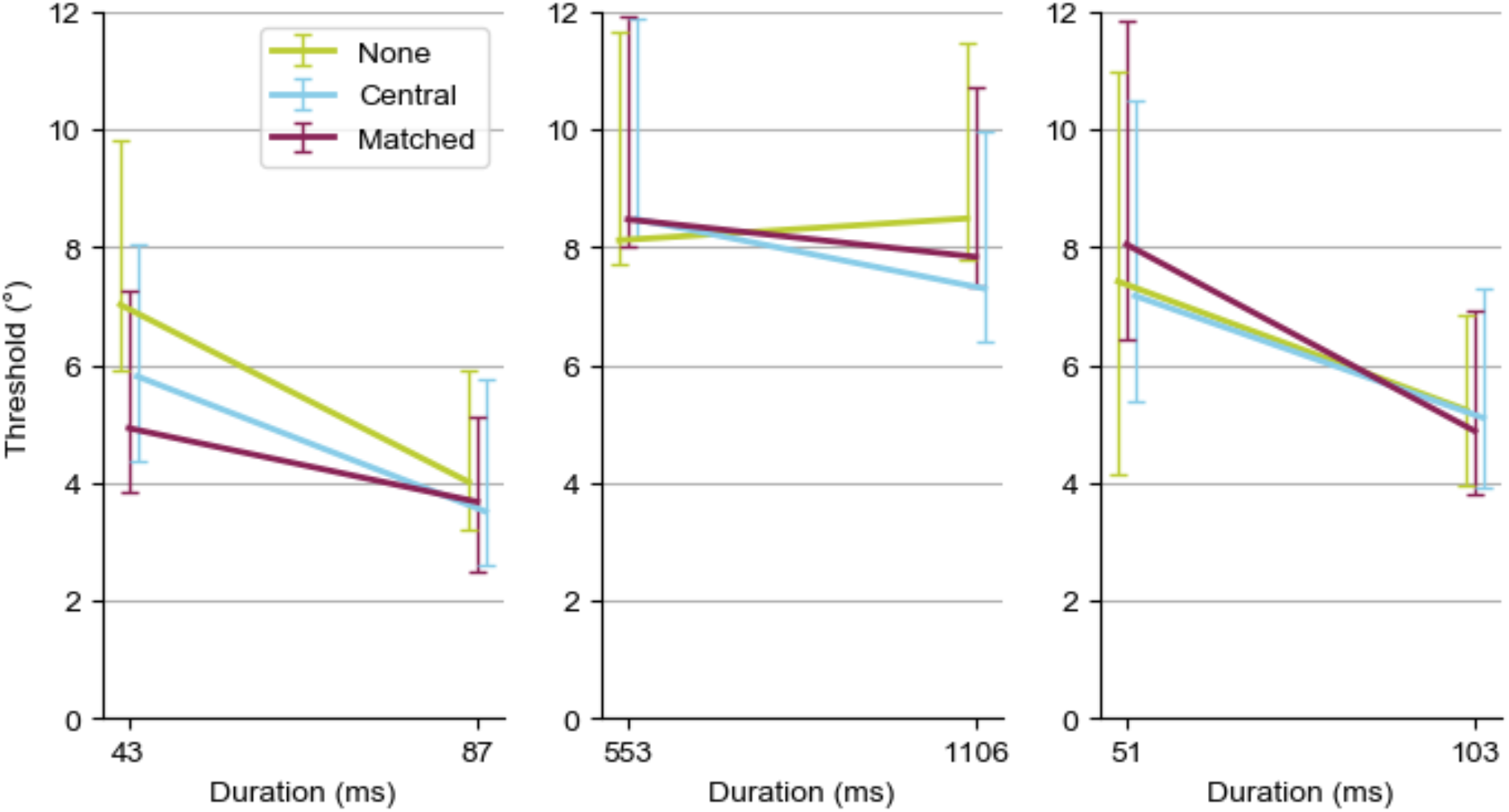
Subject thresholds averaged across all sessions (Left: Subject 1, Center: Subject 2, Right: Subject 3). Jitter along the duration axis is imposed for better visibility of each condition. Error bars show 95% Cis and are calculated by resampling daily thresholds with replacement. All subjects were able to perform the task well above chance and improved their performance at wider separations, making it possible to calculate separation thresholds for each of the six conditions (two duration x three visual). Raw performance by day and the sigmoid fits of this data are shown in Supplemental Figures 1 and 2, respectively. Each of the three subjects shows a different pattern of results. We fit separate linear mixed effects models (Supplemental Table 2–4) for each subject. The effect of separation was significant for all subjects, and the effect of duration was significant for subjects 1 and 3. Visual condition and interactions of visual condition with separation and duration were not significant predictors of behavior for any subject. This pattern can be most clearly seen in the thresholds (Supplemental Figure 3), with subject 1 and 3 showing a decrease/improvement in threshold going from the shorter to the longer duration. To summarize, none of the parameters of interest showed significant effects for any individual subject.

**Supplemental Table 2:** GLM results for Experiment 2 Subject 1.

|  | Estimate | Std Err | Z Value | P Value |
| --- | --- | --- | --- | --- |
| Vis Central vs. Matched | 0.565 | 0.596 | 0.948 | 0.343 |
| Vis None vs. Matched | -0.402 | 0.596 | -0.674 | 0.500 |
| Separation | 1.87 | 0.203 | 9.22 | $<2 \times 10^{-16}$ |
| Duration | 0.890 | 0.278 | 3.20 | 0.00136 |
| Vis Central interaction with Duration | -0.118 | 0.373 | -0.316 | 0.752 |
| Vis None interaction with Duration | 0.0594 | 0.371 | 0.160 | 0.873 |
| Vis Central interaction with Separation | -0.363 | 0.265 | -1.37 | 0.171 |
| Vis None interaction with Separation | -0.00296 | 0.269 | -0.011 | 0.991 |

**Supplemental Table 3:** GLM results for Experiment 2 Subject 2.

|  | Estimate | Std Err | Z Value | P Value |
| --- | --- | --- | --- | --- |
| Vis Central vs. Matched | 0.0428 | 0.476 | 0.090 | 0.928 |
| Vis None vs. Matched | 0.119 | 0.468 | 0.254 | 0.800 |
| Separation | 1.95 | 0.152 | 12.8 | $<2 \times 10^{-16}$ |
| Duration | 0.0335 | 0.183 | 0.183 | 0.855 |
| Vis Central interaction with Duration | 0.294 | 0.262 | 1.12 | 0.261 |
| Vis None interaction with Duration | -0.0824 | 0.257 | -0.321 | 0.749 |
| Vis Central interaction with Separation | -0.0531 | 0.216 | -0.246 | 0.806 |
| Vis None interaction with Separation | -0.0585 | 0.212 | -0.275 | 0.783 |

**Supplemental Table 4:**
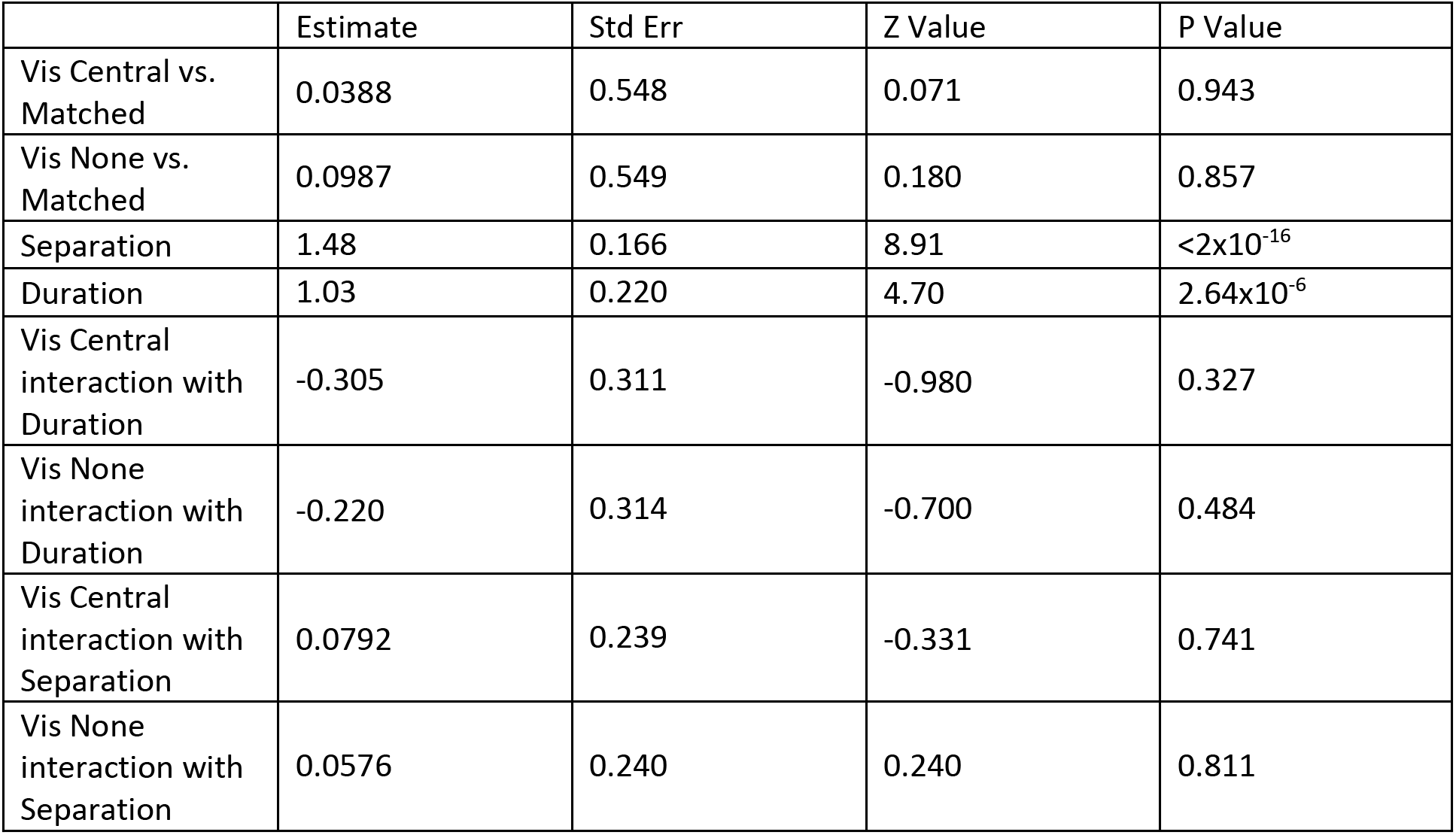
GLM results for Experiment 2 Subject 3.

## REFERENCES

Alais, D. & Burr, D. (2004) The Ventriloquist Effect Results from Near-Optimal Bimodal Integration. Current Biology, 14, 257–262.

Algazi, V.R., Duda, R.O., Thompson, D.M., & Avendano, C. (2001) The CIPIC HRTF database. In Proceedings of the 2001 IEEE Workshop on the Applications of Signal Processing to Audio and Acoustics (Cat. No.01TH8575). Presented at the 2001 IEEE Workshop on the Applications of Signal Processing to Audio and Acoustics, IEEE, New Platz, NY, USA, pp. 99–102.

Bates, D., Mächler, M., Bolker, B., & Walker, S. (2015) Fitting Linear Mixed-Effects Models Using **Ime4**. J. Stat. Soft., 67.

Blaser, E., Pylyshyn, Z.W., & Holcombe, A.O. (2000) Tracking an object through feature space. Nature, 408, 196–.

Cappelloni, M.S., Shivkumar, S., Haefner, R.M., & Maddox, R.K. (2019) Task-uninformative visual stimuli improve auditory spatial discrimination in humans but not the ideal observer. PLOS ONE, 14, e0215417.

Ernst, M.O. & Banks, M.S. (2002) Humans integrate visual and haptic information in a statistically optimal fashion. Nature, 415, 429–433.

Kaernbach, C. (1991) Simple adaptive testing with the weighted updown method. Perception & Psychophysics, 227–229.

Körding, K.P., Beierholm, U., Ma, W.J., Quartz, S., Tenenbaum, J.B., & Shams, L. (2007) Causal Inference in Multisensory Perception. PLOS ONE, 2, e943.

Larson, E. & Lee, A.K.C. (2013) The cortical dynamics underlying effective switching of auditory spatial attention. NeuroImage, 64, 365–370.

Saberi, K. (1995) Some considerations on the use of adaptive methods for estimating interaural-delay thresholds. The Journal of the Acoustical Society of America, 98, 1803–1806.

Sansone, L.G., Stanzani, R., Job, M., Battista, S., Signori, A., & Testa, M. (2021) Robustness and static-positional accuracy of the SteamVR 1.0 virtual reality tracking system. Virtual Reality,.

Shivkumar, S., Cappelloni, M.S., Maddox, R.K., & Haefner, R.M. (2022) Inferring sources of suboptimality in perceptual decision making using a causal inference task. bioRxiv, 2022.04.28.489925.

Witten, I.B. & Knudsen, E.I. (2005) Why Seeing Is Believing: Merging Auditory and Visual Worlds. Neuron, 48, 489–496.

